# Characterization of novel regulators for heat stress tolerance in tomato from Indian sub-continent

**DOI:** 10.1101/800607

**Authors:** Sonia Balyan, Sombir Rao, Sarita Jha, Chandni Bansal, Jaishri Rubina Das, Saloni Mathur

## Abstract

The footprint of tomato cultivation, a cool region crop that exhibits heat stress (HS) sensitivity, is increasing in the tropics/sub-tropics. Knowledge of novel regulatory hot-spots from varieties growing in the Indian sub-continent climatic zones could be vital for developing HS-resilient crops. Comparative transcriptome-wide signatures of a tolerant (CLN1621L) and sensitive (CA4) cultivar-pair short-listed from a pool of varieties exhibiting variable thermo-sensitivity using physiological, survival and yield-related traits revealed redundant to cultivar-specific HS-regulation with more up-regulated genes for CLN1621L than CA4. The anatgonisiticly-expressing genes include enzymes; have roles in plant defense and response to different abiotic stresses. Functional characterization of three antagonistic genes by overexpression and TRV-VIGS silencing established Solyc09g014280 (*Acylsugar acyltransferase*) and Solyc07g056570 (*Notabilis*), that are up-regulated in tolerant cultivar, as positive regulators of HS-tolerance and Solyc03g020030 (*Pin-II proteinase inhibitor*), that is down-regulated in CLN1621L, as negative regulator of thermotolerance. Transcriptional assessment of promoters of these genes by SNPs in stress-responsive *cis*-elements and promoter swapping experiments in opposite cultivar background showed inherent cultivar-specific orchestration of transcription factors in regulating transcription. Moreover, overexpression of three ethylene response transcription factors (ERF.C1/F4/F5) also improved HS-tolerance in tomato. This study identifies several novel HS-tolerance genes and provides proof of their utility in tomato-thermotolerance.

**Highlight:** Novel heat stress regulatory pathways uncovered by comparative transcriptome profiling between contrasting tomato cultivars from Indian sub-continent for improving thermotolerance. (20/30)

## Introduction

Global warming is exposing plants to unfavourable temperature fluctuations resulting in detrimental effects on crop productivity and an ultimate threat to biodiversity and food security (Battisti et al., 2009; Challinor et al., 2014). As per the 5^th^ report of Intergovernmental Panel on Climate Change (IPCC), 0.8-4.8°C rise in the global mean temperatures has been projected by the end of this century (IPCC, 2015). High temperature affects the phenological development, photosynthesis, respiration, initiation, expansion and senescence of plant organs (Wanget al., 2017). As sessile life forms, it is imperative for plants to evolve mechanisms to adapt to the thermal stress to minimise damage on its growth and reproduction. This requires dynamic reprogramming at the level of transcriptome, proteome, metabolome, lipidome and epigenome leading to the activation of complex heat stress response (HSR) (Mittler et al., 2012). It is therefore, necessary to have in-depth understanding of the plant thermotolerance mechanisms at molecular level to achieve sustainable food production. The canonical HSR involves HS sensing by Phytochrome B (Jung et al., 2016), the heat induced modulations at the level of membrane stability that activates the calcium and lipid signalling (Saidi et al., 2010), as well as, the interplay of conserved heat stress transcription factor-heat shock protein (HSF-HSP) module. Upon HS, HSFA1s, the master regulators of HSR, are relieved from the repression by HSP70/90, relocate from cytoplasm to nucleus and activate other HSFs (HSFA7s, HSFA2 and HSFBs), dehydration-responsive element-binding protein 2A (DREB2A) and multiprotein bridging factor 1C (MBF1C) (Mishra et al., 2002; Liu et al., 2011; Yoshida et al., 2011). These transcriptional regulators further control expression of several HS-inducible genes including HSFA3, nuclear transcription factors Y (NFYs) and HSPs (Schramm et al., 2008; Yoshida et al., 2008, 2011; Chen et al., 2010; Sato et al., 2014). Another critical regulator of thermotolerance is the C-REPEAT BINDING FACTOR2 (RCF2) regulated NAC019 that acts as an upstream regulator of HSFA1b, HSFA6b, HSFA7a and HSFC1 (Guan et al., 2014). Several HSF-target modules like HSFA4a-APX1 and HSFA5-HSPs, act independent of HSFA1s pathway (Liu et al., 2006; Von Koskull-Doring et al., 2007). Apart from these canonical HSR circuits, PIF4 regulatory and associated signalling networks control plant thermo-morphogenesis (Koini et al., 2009; Stavang et al., 2009; Oh et al., 2012; Proveniers and Van Zanten, 2013). The significance of post-translational regulation in HS-response is also well documented in plant HSR (Agarwal et al., 2007; Miller et al., 2010; Hahn et al., 2011; Vainonen et al., 2012; Mizoi et al., 2013; Sato et al., 2016).

Tomato is the second most cultivated vegetable in the world. HS leads to reduced tomato yield by affecting its vegetative as well as reproductive development (Sato et al., 2000; Pressman et al., 2002). Transcript profiling using cDNA-AFLP (Bita et al., 2011) and microarray analysis (Frank et al., 2009) as well as proteomic analyses (Mazzeo et al., 2018) of microspores have shown active involvement of HSPs, ROS scavengers, hormones, amino acid metabolism and nitrogen assimilation in tomato HS-response. Heat inducible tomato HSP21 protects PSII from the HS-induced oxidative stress and fruit ripening (Neta-Sharir et al., 2005). Tomato HSFA2 is highly up-regulated during HS that in turn forms a hetero-oligomeric transcriptional complex with HSFA1. In addition, HSP17.4-CII acts as a co-repressor of HSFA2 (Port et al., 2004). Over-expression of Arabidopsis Receptor-like kinase ERECTA (ER) leads to enhanced HS tolerance in Arabidopsis, rice and tomato with increased biomass (Shen et al., 2015). Till now the knowledge about the effect of HS on tomato leaf exploiting the comparative transcriptomics of contrasting cultivars in response to heat is not well understood. Tomato is increasingly becoming a cash crop in regions of higher temperatures (the tropics and sub-tropics) than the optimum 26/20 °C during the photoperiod/dark period (Srivastava et al., 2012). One way for developing HS-resilient crops is to exploit the naturally evolved mechanistic signalling frameworks in cultivars exhibiting contrasting response to HS (Challinor et al., 2014). To uncover the HS-responsive molecular mechanisms in tomato using naturally occurring cultivars growing in the sub-tropical climatic conditions of the Indian sub-continent, we first evaluated nine tolerant/sensitive tomato cultivars under HS to identify the best contrasting pair viz., CLN1621L (tolerant) and CA4 (sensitive). Comparative analysis of the heat-responsive transcriptomic signatures of leaves highlighted conserved to genotype-specific reprogramming at transcriptional levels. Functional assessment by silencing (Virus-induced gene silencing, VIGS) and transient over-expression of three previously uncharacterised genes, that exhibit antagonistic expression in the contrasting varieties under HS, established their roles as positive/negative regulators of HS-tolerance. Promoter:reporter expression analysis highlights that cultivar-dependent transcription factors pool governs the opposite expression of these genes. The study provides a collection of heat as well as cultivar-specific genes for genetic engineering of tomato for thermotolerance.

## Materials and Methods

### Screening of tomato cultivars for HS tolerance

Seeds of tomato cultivars were obtained from Asian Vegetable Research and Development Centre (AVRDC), Taiwan (CLN1621L and CA4); Indian Agricultural Research Institute (IARI), New Delhi, India (Pusa-120, Pusa Rohini, Pusa Sadabahar and Pusa Ruby); CCS, Haryana Agricultural University (CCS,HAU), Hisar, India (Hisar Arun) and Indian Institute of Horticultural Research (IIHR), Bengaluru, India (IIHR-2274 and IIHR-2201). To mimic the natural stress conditions, all the cultivars were grown in field under natural condition for two years and two seasons each at experimental fields of National Institute of Plant Genome Research (NIPGR), New Delhi, India, one in optimum growing conditions and second when temperatures were high followed by their extensive physiological and yield based analysis. The detailed methodology of screening of tomato cultivars is described in Supplementary methods S1.

### RNA-Seq Analysis

The paired-end libraries of leaf of tolerant and sensitive cultivars were constructed and sequenced on illumina platform with three biological replicates from control and HS conditions each. The raw data was analysed using the RNA-Seq tool of CLC genomics workbench. For further details on the transcriptome experiment and data analysis for differential gene expression, refer to Supplementary Methods S1.

### Expression analysis

The expression analysis was performed as per Paul et al. (2016). The relative fold change was calculated by following the ddCT method using tomato actin gene as endogenous control. The sequence of primers used is provided in Supplementary Table S1.

### Bioinformatics tools used

The GO-enrichment analysis was performed using the “Gene list analysis” tool of PANTHER classification system (www.pantherdb.org, version 14.1) following the statistical over-representation test using default settings (Bonferroni correction P-value ≤0.05; Fold enrichment ≥2). The KEGG pathway enrichment analysis was done using ShinyGO version 0.60 (P-value ≤0.05). The 1kb upstream regulatory region of genes were retrieved from Sol Genomics Network (https://solgenomics.net) and analysed for cis-regulatory elements using the Plant Promoter Analysis Navigator (PlantPAN; http://plantpan2.itps.ncku.edu.tw/) using multiple promoter analysis tool.

### In-planta transient assays

Transient silencing assays following the virus induced gene silencing approach (VIGS) was performed as described by Muthappa et al., 2014. The transient overexpression of selected genes followed by thermotolerance assays were standardized for different tomato cultivars by following the methodology adopted by Queitsch et al. (2000) with modifications. For cloning of promoter, 1.5kb region upstream of start sites of genes were amplified from genomic DNA of contrasting cultivars and cloned into pBI101 in fusion with GUS. The sequence of primers is present in Supplementary Table S1. For detailed methods refer the supplementary methods S1.

### Gas-exchange measurements

Tomato leaf gas-exchange measurements, including water use efficiency (WUEi), net photosynthetic rate (A), transpiration rate (E) and stomatal conductance (Gs) were measured simultaneously by using a portable Licor 6400 photosynthesis system (LI-6400, Li-Cor Inc., Lincoln NE, USA). For all measurements, sixth and seventh fully expanded leaf of 6 plants for each construct were used and the experiment was repeated twice with similar parameters. The measurement conditions were as follows: leaf temperature 27°C, leaf–air vapour pressure deficit 1.5±0.5 kPa, photosynthetic photon flux 300 μmol m^−2^ s^−1^, relative air humidity 70% and ambient CO_2_ concentration 400±5 μmolmol^−1^.WUEi was calculated as using following formula A/E.

### Histochemical detection of H_2_O_2_ and cell death

For estimating the hydrogen peroxide (H_2_O_2_) levels using 3,3’-diaminobenzidine (DAB) and cell death using trypan blue, 6th and 7th leaf from the 6-week-old TRV and TRV-gene plants were used immediately after HS (45°C for 4.5 h). DAB staining was performed as previously described (Daudi et al., 2012). The viability of leaf cells was measured using the trypan blue exclusion method (Koch and Slusarenko 1990). Experiment was repeated twice with similar parameters with sample size of 6 plants per replicate.

## Results and Discussion

### CLN is heat stress tolerant and CA4 is sensitive tomato cultivar

The genomic variations between different cultivars within the same species are often armed with specific mechanisms to tolerate stress at morphological, physiological and molecular levels (Huang and Gao, 1999). Tomato production has considerably increased in the tropical and sub-tropical regions (Nicola et al., 2009; Srivastava et al., 2012). To exploit the naturally occurring variability between tomato cultivars growing under Indian sub-continent climatic conditions for enhancing thermotolerance, we performed comparative assessment of nine tomato cultivars known to be either HS tolerant or sensitive to short-list best contrasting pair. These cultivars included five tolerant [CLN1621L (henceforth referred to as CLN), IIHR2201, IIHR2274, Pusa Sadabahar and Hisar Arun] and four sensitive cultivars (CA4, Pusa Ruby, Pusa Rohini, and Pusa120) (Chavan et al., 2009; Kartikeya et al., 2012; Meena et al., 2015; Sangu et al., 2015). The tolerance of cultivars to HS was judged on parameters adopted by several scientific groups to screen tomato genotypes (Baki et al., 1991; Baki and Stommel, 1995). These include survival assays at seedling stage (Supplementary Figure S2A) and at 1-month-old stage (Supplementary Figure S2B), physiological assays like proline content, RWC, and EL (Supplementary Figure S2C-E) as well as yield-related traits like percent fruit set per plant and total fruit yield (Supplementary Figure S2F-G). Higher values for all the above parameters and lower for EL are hallmarks of better thermotolerance. The cultivars were ranked for their performance by assaying all the above parameters as percent increase/decrease under HS relative to control conditions. CLN displayed highest ranking among the cultivars for all the above parameters except EL (Figure 1A and Supplementary Figure S2E). CA4 on the other hand had highest EL and obtained lowest rank for all other parameters except proline content where Pusa Ruby had least value (Figure 1A; Supplementary Figure S2C). Of note, percent survival of 1-month-old CLN plants was more than 80% whereas it was mere 11% in CA4. Fruit set was also higher in CLN (81%) as compared to CA4 (27%), that in turn reflected by nearly 50% yield being maintained in CLN but dropped to 6% in CA4 (Supplementary Figure S2F-G). Hierarchical clustering clearly divided the cultivars into four clades based on their HS response viz. (I) most tolerant (CLN), (II) moderately tolerant (Pusa Sadabahar, IIHR2274, Hisar Arun and IIHR2201), (III) sensitive (Pusa Ruby, Pusa-120 and Pusa Rohini) and (IV) highly sensitive (CA4) (Figure 1A). This was further strengthened by PCA of all the above factors that showed the separation of each cultivar on PC1 (82.8%) based on the performance under HS, placing CLN and CA4 on the two extremities (Figure 1B). Thus, identifying them as the most tolerant and sensitive cultivars to HS, respectively. CLN is reported to be tolerant to salt stress (Saeed et al., 2011) and TMV infection (https://avrdc.org). CLN has been shown to have higher plant vigour and pollen viability as compared to sensitive CA4 (Sangu et al., 2015). In addition, CLN maintains highest percentage of seeded fruits and lowest percentage of parthenocarpic fruits under HS as compared to several other cultivars (Comlekcioglu and Kemel-Soylu, 2010). The CLN-CA4 pair has been used as donor parents for generating mapping populations towards HS (https://avrdc.org). Thus, decoding of transcriptome of these two will yield critical insights into HS response and provide valuable resource for future research on thermotolerance improvement in tomato.

**Figure 1.**
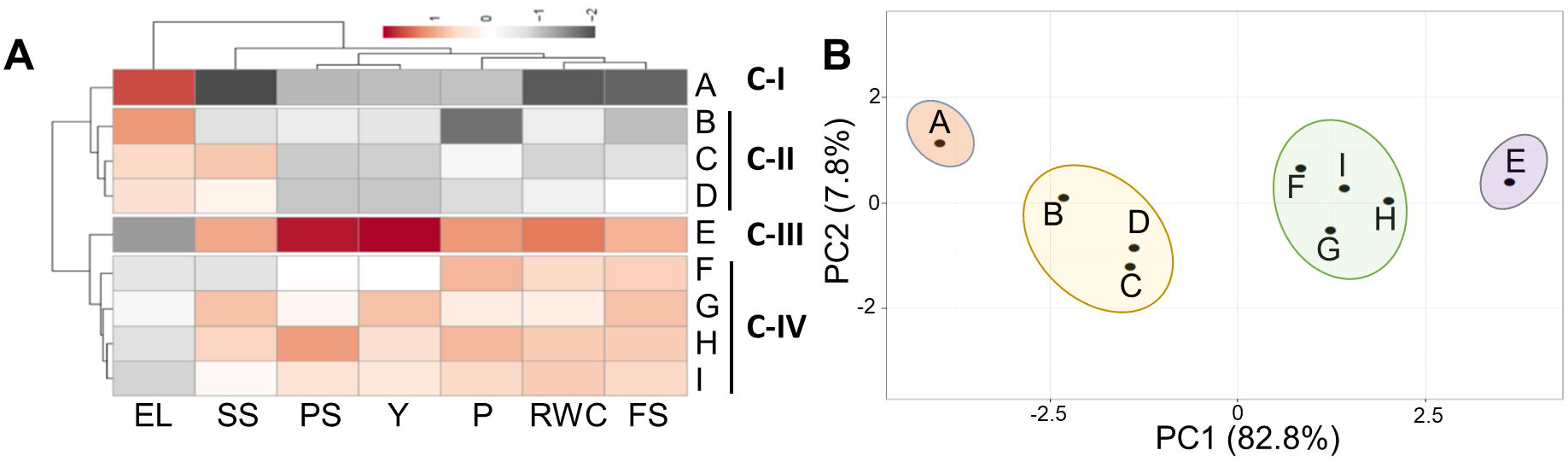
Identification of heat stress tolerant and sensitive tomato cultivars. (A) Nine tomato cultivars with variable heat stress tolerance were screened to select best contrasting pair to heat stress response by assessing physiological, biochemical and yield related traits. Based on the performance of each cultivar, the percentage value of each trait was calculated under heat stress relative to respective controls which was set as 100%. All these traits were analysed by clustering that divided them in four classes I to IV. (B) Principal component analysis of the nine cultivars on above parameters using ClustVis. Clustering was performed using Manhattan distance and complete linkage. Unit variance scaling was applied to rows and Singular Value Decomposition (SVD) with imputation was used to calculate principal components. Cultivars are represented by alphabets A to I. A: CA4; B: Pusa Ruby; C: Pusa-120; D: Pusa Rohini; E: CLN1621L; F: Hisar Arun; G: IIHR2274; H: Pusa Sadabahar; I: IIHR220. EL: Electrolytic Leakage; SS: 5-day-old seedling survival; PS: 1-month-old plant survival; Y: Yield; P: Proline content; RWC: Relative water content; FS: Percent fruit set/plant.

### Distinct transcriptome signatures between two cultivars

Response to abiotic stress is a complex trait involving a network of genomic elements, therefore, transcriptome-wide comparative landscape of the selected tolerant-sensitive cultivar-pair should shed insights into regulatory ‘hotspots’ to understand thermotolerance mechanism. In nature, plants protect themselves from severe damage to extreme HS by acclimating themselves to sub-lethal temperature stresses. To mimic similar conditions in the study, the plants were first acclimated to HS by exposing them to milder HS before exposing to harsh temperatures (see material and methods). More than 55 million good quality trimmed reads each for control and HS-response were obtained, the paired-end datasets exhibited 95-97% mapping on tomato transcripts in different datasets (Supplementary Table S2). Out of 35,768 genic loci (ITAG release 3.2) of tomato, 26,631 genes were detected in present study (Supplementary Table S3). The remaining genes are either very low expressing or could be other development stage/stress-specific. PCA of transcriptome samples showed very good correlation between the biological repeat datasets and demonstrated a clear transcriptomic disparity between the two cultivars as well as distinctly separated the stress and control samples (Supplementary Figure S3A). This was also substantiated by hierarchical clustering of the control and heat transcriptomic datasets; they being divided into two clusters (Supplementary Figure S3B). The hierarchical clustering of 35,768 genes showed core to HS-specific regulation of several gene clusters (Supplementary Figure S3C).

### Universal to cultivar-specific regulation to HS

We next examined only that fraction of transcriptome that was significantly modulated (false discovery rate FDR p-value ≤ 0.05 and fold change ≥ 2 and ≤-2 for up- and down-regulated genes, respectively) in leaf under HS as depicted by volcano plots (Supplementary Figure S3D). A total of 6954 differentially expressing genes (DEGs) were identified between both cultivars (Figure 2A). This number is the highest number of DEGs so far in tomato and the first for leaf utilising contrasting cultivars (Bita et al., 2011; Fragkostefanakis et al., 2015; Fragkostefanakis et al., 2016; Keller et al., 2018). A slightly higher number of HS-responsive genes are present in the HS-tolerant cultivar (5239 vs. 5138 in CLN and CA4, respectively), moreover the up-regulated events are more in tolerant cultivar {2580 (CLN) and 2496 (CA4)} while almost equal number of down-regulated genes are present in both cultivars {2659(CLN) and 2642 (CA4)} (Figure 2A). A high magnitude of up-regulation has also been reported in anthers of heat tolerant (Heat Set 1) tomato genotype in comparison to sensitive genotype, Falcorosso (Bita et al., 2011). Surprisingly, only 21 genes showed an opposite (antagonistic) HS-regulation between CLN and CA4 (Figure 2A-B); 5 genes viz., Solyc03g020030 (PIN-type-II proteinase inhibitor 69), Solyc09g084450 (proteinase inhibitor I), Solyc12g013700 (Stem-specific protein tsjt1), Solyc01g079530 (RING/FYVE/PHD zinc finger superfamily protein) and Solyc12g042500 (Gibberellin-regulated family protein) are down-regulated in CLN but up-regulated in CA4 and 16 genes viz., Solyc01g065530 (Protein COBRA), Solyc01g081250 (Glutathione s-transferase), Solyc01g087020 (Transmembrane protein), Solyc01g095140 (Late embryogenesis abundant protein), Solyc01g107780 (Glycosyltransferase), Solyc03g116890 (WRKY transcription factor 39), Soly05g052670,Solyc05g052680 andSolyc09g014280 (all threeHXXXD-type acyl-transferase family proteins), Solyc06g049070 (Nucleotide/sugar transporter family protein), Solyc07g008103 (Blue copper protein), Solyc07g056570 (Notabilis), Solyc07g064410 (Kua-ubiquitin conjugating enzyme hybrid), Solyc09g075920 (Serine/threonine-protein kinase), Solyc09g092500 (Glycosyltransferase) and Solyc10g081570 (Marmande) are up-regulated in CLN but down-regulated in CA4. Gene ontology (GO) enrichment of these antagonist genes highlights the significant enrichment of negative regulation of endopeptidase, peptidase and proteolysis, response to water, pollination and transferase activity (Figure 2C). Among these, five (the two proteinase inhibitors and three HXXXD-type acyl transferases) have roles in plant defense. The qPCR validation of 10 genes selected from the above clusters confirms the opposite HS regulation in CA4 and CLN (Figure 2D).

**Figure 2.**
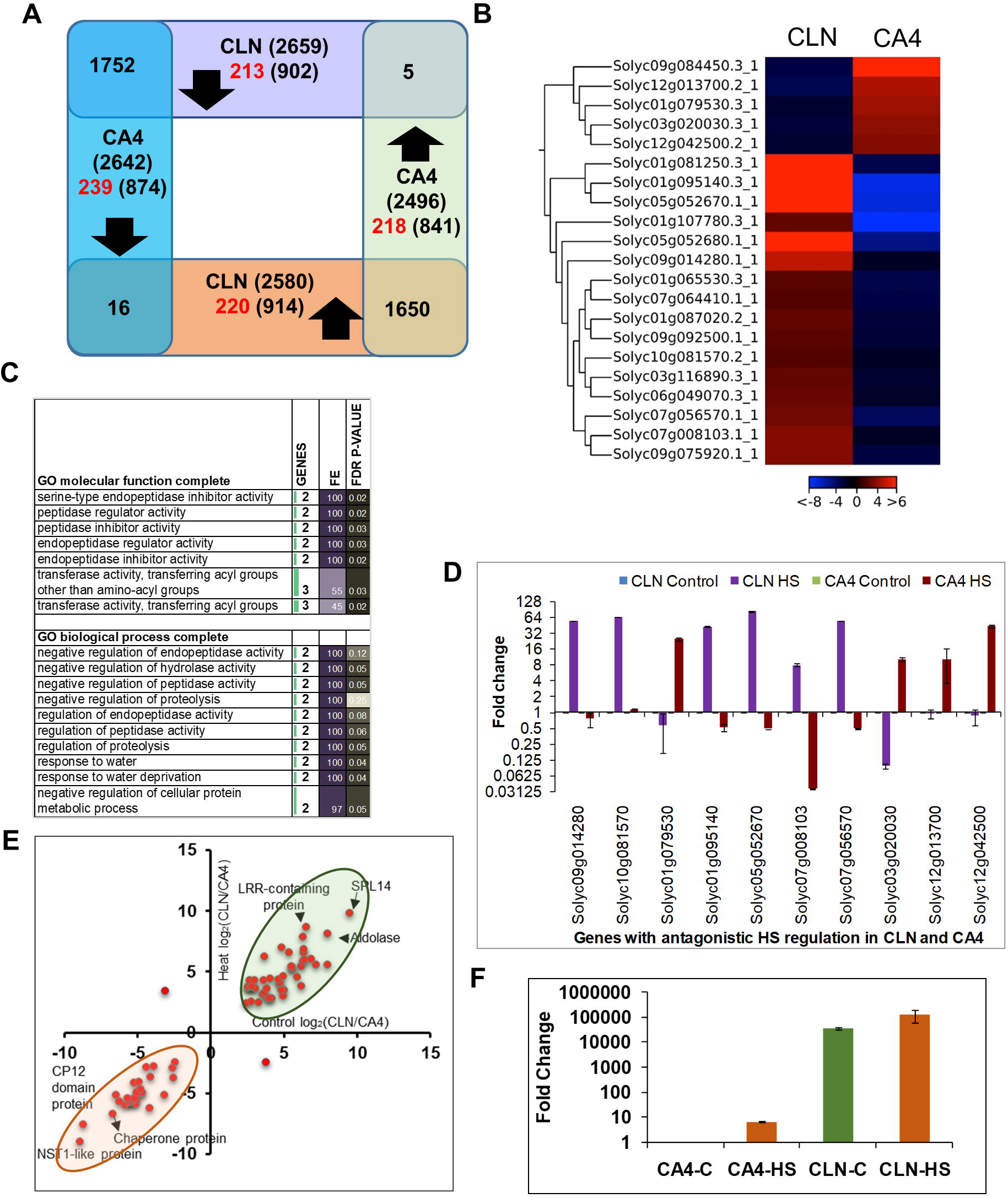
Heat stress responsive transcriptional modulations in leaf of tolerant, CLN and sensitive, CA4 tomato cultivars. **(A)** The venn diagram depicts the comparison of heat response of CLN and CA4 leaf transcriptome. The genes with at least 2 fold up- or down-regulation with FDR p-value of ≤0.05 were considered. The genes with strict cultivar-specific HS response are written in red. **(B)** The expression profile of cluster of genes with opposite heat-mediated regulation in leaf of CLN and CA4 under control and heat conditions. The heat map shows the fold up/down regulation of genes under HS in CLN and CA4. **(C)** GO enrichment analysis of molecular function and biological process of the genes with inverse HS regulation in tolerant and sensitive cultivar following the PANTHER Overrepresentation Test. **(D)** The qPCR validation of selected genes with antagonistic HS-response between CLN and CA4 in leaves. **(E)** The scatter plot showing the genes with ≥ 5 fold higher expression in control as well as in heat conditions in CLN (green) and in CA4 (orange). Details of all these genes is provided in Supplementary Table S5. **(F)** qRT-PCR validation of *SPL 14* gene in CA4 and CLN leaf under control and HS conditions, normalized with CA4 control. In qPCR, two biological with three technical repeats and actin as endogenouse control was used.

Around ~49% (3402 out of 6954) HS-responsive genes followed conserved HS-regulated gene expression; 1752 DEGs being up- and 1650 DEGs being down-regulated in both cultivars (Figure 2A), signifying a core universal HS response. GO enrichment analysis suggests the involvement of up-regulated genes in cellular response to nutrient starvation specifically phosphate starvation in addition to protein folding (Supplementary Figure S4A-B). Further proteins belonging to chaperones, winged helix TFs, mRNA splicing and processing factors are significantly enriched among the conserved up-regulated genes (Supplementary Figure S4C). KEGG pathways like protein processing in ER, spliceosome and ubiquitin mediated proteolysis were enriched in conserved up-regulated genes (Supplementary Table S4). Many recent reports on stress-responsive regulation of spliceosome per se, as well as, crucial role of alternatively spliced transcripts in response to abiotic stresses including HS have been reported in plants (Deng et al., 2011; Seo et al., 2012, 2013; Carrasco-López et al., 2017; Calixto et al., 2018; Filichkin et al., 2018; Laloum et al., 2018; Huertas et al., 2019). The unfolded protein response (UPR) is a conserved response that is elicited by ER stress and is known to protect plants from adverse environmental stresses (Wan and Jiang, 2016; Li et al., 2018; Park and Park, 2019). On the other hand, wide range of biological processes and metabolic pathways belonging to various protein classes were down-regulated in both the cultivars (Supplementary Figure S5-7; Table S4). We find that 26 genes are highly up-regulated (≥100-folds) in both cultivars but none followed similar folds’ down-regulation in both (Supplementary Table S3). The highly expressed genes (>1000 fold) in CLN and CA4 belong to the HSP protein family (Solyc03g113930.2.1, Solyc01g102960.3.1, Solyc03g082420.3.1, Solyc11g020330.1.1) and a serine carboxypeptidase (Solyc04g077640.3.1).

Nearly 51% (3531) of HS-responsive genes follow cultivar-biased stress regulation, i.e., those DEGs which are significantly up- or down-regulated in only one cultivar but not in other. These include 1816 (914 up, 902 down) DEGs in CLN and 1715 (841 up, 874 down) DEGs in CA4 (Figure 2A). To further discriminate the true cultivar biased HS-responsive DEGs, the above genes were filtered out by adopting strict parameters (See Material and methods). As a result, 218 up and 239 downregulated genes showed CA4-specific HS regulation while 220 up and 213 downregulated genes followed CLN-specific HS response. (Figure 2A; Supplementary Table S3). Genes exhibiting the CLN specific down-regulation were enriched in terms associated with photosynthesis in biological process and cellular component category while glycopeptide alpha-N-acetylgalactosaminidase activity, cystein synthase activity, chlorophyll binding, ATPase activity were enriched among the molecular function (Supplemental figure S8). For genes exhibiting the CLN-specific upregulation, there was no significant enrichment. While, there was significant enrichment of translation, ribosomal proteins, ribosome, structural constituents of ribosomes etc in CA4-specific upregulatated and role of calcium mediated signalling and GTPase activity were highlighted among the CA4-specific downregulated genes (Supplemental figure S9-10).

In addition, some metabolic pathways were enriched specifically in up-regulated genes in CLN only *viz.* MAPK signalling pathway, circadian rhythm, plant hormone signal transduction, sulphur metabolism and glycerolipid metabolism (Supplementary table S4). Tomato, MPK1 regulates thermotolerance via SlSPRH1 involved in antioxidant defense (Ding et al., 2018). Phosphorylation is critical regulatory mechanism for several HSFs (Evrard et al., 2013; Link et al., 2002). The circadian clock genes sustain healthy and accurate timing over an array of physiological temperatures in different plant species. HSFB2b-mediated transcriptional repression of PRR7 is known to direct HS responses of the circadian clock in *Arabidopsis* (Kolmos et al., 2014). Also, the central component of circadian clock, ZEITLUPE and HSP90 have been found to be critical for the maintenance of thermo-responsive protein quality (Gil et al., 2017;Gil and Park, 2017). Our data contributes to identification of HS-responsive signature genes which can be exploited for thermotolerance augmentation in tomato and other crop plants.

### Several genes follow cultivar-specific expression pattern

Further, we assessed whether there is any inherent genotype-based difference in the relative transcript abundance between the contrasting varieties under control or HS conditions. For this analysis we used more stringent criteria of fold change of ≥5 with FDR p-value correction of ≤0.05 and found 86 and 50 highly abundant transcripts in CLN and CA4, respectively under control conditions while 130 CA4-abundant and 231 CLN-abundant in HS conditions (Supplementary Table S5). Interestingly a cluster of 29 genes maintained high expression in CA4 under both control as well as in HS (Figure 2E, orange) while 46 such genes were identified in CLN (Figure 2E, green). Among these, Solyc04g081710 (SPL14) transcripts are >700-fold more abundant in CLN as compared to CA4 in control conditions (Figure 2E, Supplementary Table S5), this trend was also confirmed by qRT_PCR (Figure 2F). In addition, transcripts of an aldolase gene (Solyc04g081600) and DUF241 domain containing protein (Solyc02g083050) are present >250-fold more in CLN in control conditions in comparison to CA4 (Figure 2E). Three genes, Solyc06g036250 (stress response NST1-like protein), Solyc04g082910 (CP12 domain-containing family protein), Solyc09g011560 (glutathione S-transferase) and Solyc09g018960 (Chaperone Dna-J) exhibited strong expression (FC>100) in CA4 in both control and stress conditions (Figure 2E, Supplementary Table S5).

### Distinct modulation of transcription factors in CLN and CA4 under heat stress

Gene regulation by TFs acts as focal nodes in all molecular networks. We predicted 1963 tomato TFs belonging to 58 families, using the TF prediction server of PlantTFDB (http://planttfdb.cbi.pku.edu.cn/prediction.php) with protein sequences of 35768 genes (ITAG version 3.2) as query (Supplementary Table S6). Out of these, 402 TFs belonging to 46 families were found to be heat-responsive in at least one cultivar. In CLN and CA4, 314 (152 down/162 up) and 285 (152 down/133 up) TFs were significantly differentially regulated, respectively (Figure 3A, Supplementary Table S6). Similar HS-response was evident for 201 (100 down/101 up) TFs in both cultivars (Figure 3A). Thirty-seven TFs (29 and 16 in CLN and CA4, respectively) exhibited very strong heat induction (FC>10). Thirty TFs followed conserved high (FC>5) up-regulation in both the cultivars while 26 followed high (FC <-5) down-regulation in both the cultivars. These highly transcribed TFs include members belonging to ERF (5 members), CO-like (3 members), HSFs (4 TFs); 4 members each of MYB and C3H, C2C2-CO-like andNAC (2) and 1 each of B3, bHLH, bZIP, C2H2, Dof1, FAR1, HD-ZIP1 and NF-YB D (Figure 3B). Of these, 3 TFs viz., HSFA2 (Solyc08g062960) and dehydration responsive element binding (DREB) TF (Solyc05g052410) were more than 100-folds up-regulated in both cultivars. The plot of above 30 TFs clearly revealed that the heat tolerant CLN showed higher magnitude of HS–mediated induction in majority of the TFs and common key TF pool were activated upon HS in both cultivars (Figure 3B). Moreover, 26 TFs showed HS-mediated strong repression (FC≤ 5) in both cultivar (Figure 3C). While Solyc08g008280 (WRKY-53) was the top repressed candidates in CLN, Solyc10g005080 (Late elongated hypocotyl) was highly down-regulated TF in CA4 under HS (Figure 3C). In addition, other TFs exhibiting strong repression belong to TF families bHLH (4), bZIP (2), C2H2 (3), MYB (3), TCP (2), WRKY (2) and YABBY (Figure 3C and Supplementary Table S6). The highly HS-responsive TF families with more than 20 of its members significantly modulated (up- and down-) at transcript level include bHLH, bZIP, C2H2, C3H, ERF, GRAS, HD-ZIP, HSF, MYB, NAC and WRKY (Figure 3D). Further, we scanned for those TF families that had at least 2 differentially regulated members in response to HS and also had ≥80% of its members exhibiting either up- or down-regulation. Using these criteria, members belonging to TF families B3, C3H, HSF, NAC and Trihelix were majorly induced in response to HS while ARF, Dof, MADS, TCP and YABBY were primarily repressed in response to HS (Figure 3D). Thermotolerance in transgenic systems have been achieved by a C2H2 zinc finger TF, ZAT12/B. carinata (Shah et al., 2013), NAC019 (Guan et al., 2014), bZIP28 (Srivastava et al., 2014) and PIF4 (a bHLH transcription factor) that acts as a central regulator of plant thermo-sensory response in Arabidopsis (Gangappa et al., 2017; Hwang et al., 2017).

**Figure 3.**
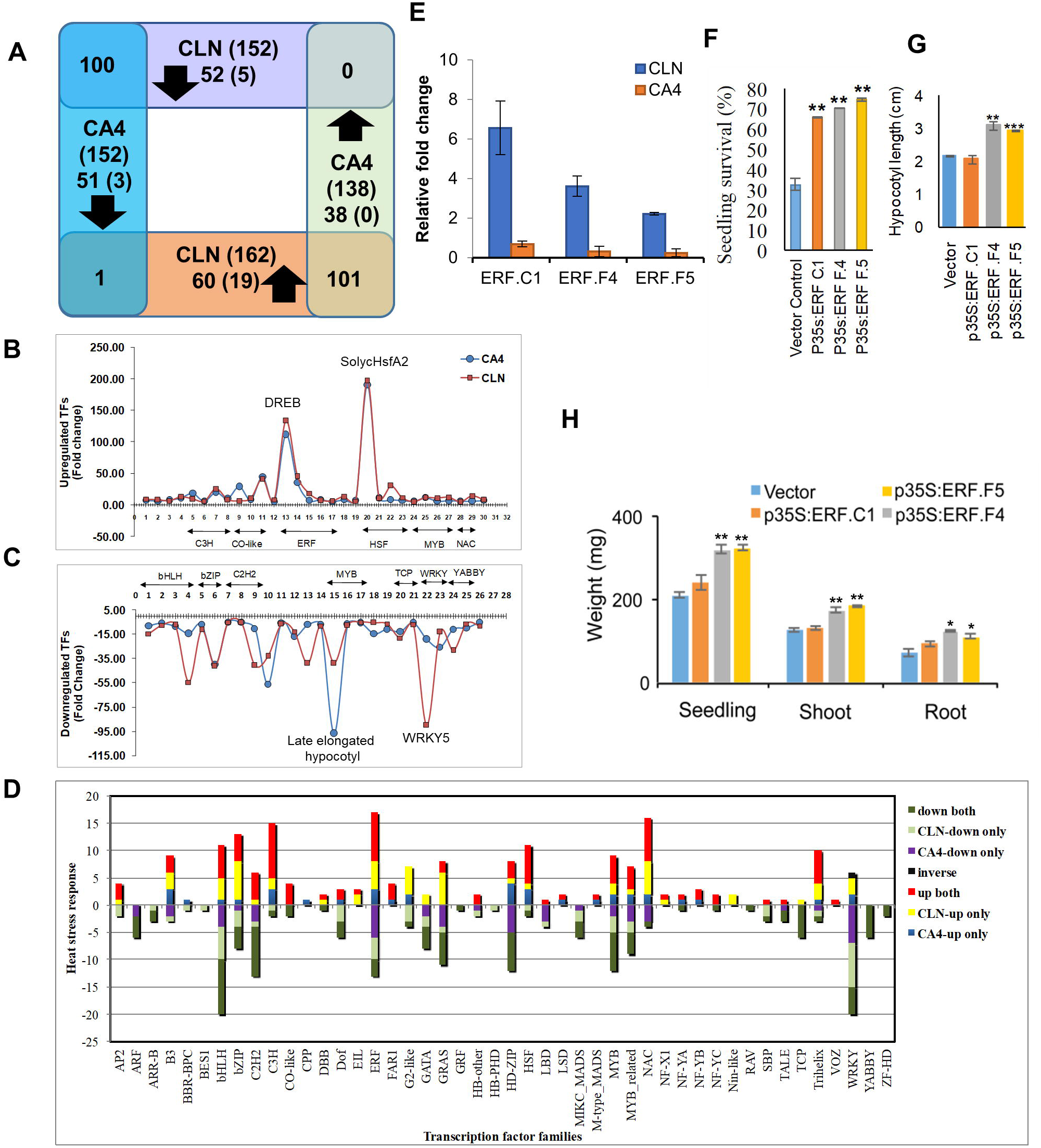
Heat stress mediated regulation of transcription factors in leaf of contrasting cultivars. **(A)** Venn diagram showing the common to specific heat responsiveness of transcription factors in leaf of tolerant, CLN and sensitive CA4. **(B-C)** The plots showing the highly up- (B, FC≥5; FDR ≤0.05) and down-regulated (C, FC ≤-5; FDR ≤0.05) transcription factors in CLN and CA4 in response to heat stress. **(D)** The comparison of cultivar specific heat response of various transcription factor families in leaf of tomato. **(E)** Expression analysis of ERFs: ERF.C1, -.D2, -.F4 and -.F5 in response to HS in leaf of CLN (tolerant) and CA4 (sensitive) cultivar by qRT-PCR. (F-G) Percent seedling survival after HS for vector control and different p35S:ERFs overexpressing seedlings and comparative morphological analysis using hypocotyl length **(G)**, seedling weight, shoot weight, root weight **(H).** The data was calculated 4-6 days post heat stress imposition from four biological sets of 70 seedlings for each gene. Error bars denote standard error. *p<0.05, **p<0.01 and ***p<0.001.

### HSF-/ERF-mediated HS regulation of CLN and CA4

The most studied and well established TF family in response to HS is the HSF family. Seven tomato HSFs (A1b, A2, A3, A4c, A5, A7 and C1), exhibited HS-mediated up-regulation in both the cultivars with highest up-regulation for HSFA2 (Supplementary Table S6). We noted that the repressor class B HSF members namely, HSFB1, B2a and B2b are up-regulated only in the sensitive CA4. Whether this has any direct correlation with CA4 being HS sensitive needs to be further evaluated. HSFB1 is endowed with both co-activator as well as repressor functions in tomato, while it acts as repressor in Arabidopsis (Fragkostefanakis et al., 2019; Bharti et al., 2004; Ikeda et al., 2011). The Arabidopsis *hsfb1-hsfb2b* double knockout mutant demonstrates higher acquired thermotolerance (Kumar et al., 2009; Ikeda et al., 2011). Also, members of the activator class A viz., HSFA6a was up-regulated and HSFA4b was down-regulated under HS in CLN only (Supplementary Table S6). AtHSFA6a is induced in response to ABA, salt and drought and is responsible for the transcriptional activation of stress responsive gene DREB2a via ABA-dependent signalling pathway (Hwang et al., 2013, 2017). Further, we investigated the ERF family that showed highest number of HS-responsive TFs and includes AP2, DREBs and ERFs (Sakuma et al., 2002). Various ERFs are reported to mediate development and stress response but only few address their regulatory role in thermotolerance (Licausi et al., 2013; Klay et al., 2014, 2018; Muller and Munne-Bosch 2015). Out of 147 ERFs reported in tomato 30 members followed differential HS response in CLN and CA4 (Supplementary Table S6). Of these, 12 ERFs were similarly regulated (9 up and 3 down) in both cultivars while others were regulated in cultivar-biased manner. We selected 4 previously uncharacterized ERFs (ERF.F4, ERF.F5, and ERF.C1) in HS-response based on the observation that all maintain high levels in CLN and low levels in CA4 under HS as validated by qRT-PCR also (Figure 3E). The above ERFs reported in fruit development (Di Metteo et al., 2013; Hao et al., 2015) in tomato and their role in thermotolerance was still lacking. Functional characterization of the above three ERFs by overexpression significantly enhanced (more than doubled) the tomato seedling survival under HS as compared to vector control plants (Figure 3F) with highest survival percentage for ERF.F5. Seedlings overexpressing ERF.F4 and F5 exhibited higher hypocotyl length as well asshoot-, root- and total seedling weight as compared to vector control plants (Figure 3G-H). Our data on TF regulation dynamics highlights role of TF families other than the undisputed relevance of HSFs in HS response. These TFs could be promising candidates for enhancing tomato thermotolerance as we demonstrated for the three ERFs that function as positive regulators of HS response.

### Role of HSPs in governing thermotolerance in tolerant-sensitive pair

The canonical plant HSR involves the co-ordinated interplay of TFs and HSPs (Frank et al., 2009; Ohama et al., 2017). A compendium of tomato HSPs was made from SGN database and literature (Paul et al., 2016; Yu et al., 2016). Out of 286 tomato HSPs belonging to various classes (Supplementary Table S7), 102 HSPs were significantly differentially regulated in either CLN or CA4. Of these,77 (69 up and 16 down) and 80 (70 up and 19 down) showed differential regulation in CLN and CA4, respectively upon HS. Since HSP class forms the heart of HS response (Queitsch et al., 2000; Su and Li 2008, Yamada et al., 2007, Wang et al., 2004), as expected majority of these HSPs were up- (60) whereas the expression of only 12 HSFs was repressed in both CLN and CA4.13 HSPs were more than 100-folds up-regulated in both varieties, of these Solyc03g113930 mRNAs rise phenomenally upon HS, increasing by ~15,000-folds and the HSFA1a/B1 regulated small HSP, Hsp21.5-ER/ Solyc11g020330 (Fragkostefanakis et al., 2019) was induced more than ~1200-folds in CLN. Several HSPs followed strict cultivar specific HS regulation. ER-resident HSP70 (Solyc06g052050) that is reported to be down-regulated in the sensitive cultivar Moneymaker under HS (Fragkostefanakis et al., 2015), is up-regulated by 16-folds in tolerant CLN. These HSPs could be critical regulators of tomato heat tolerance mechanism.

### Knocking down ASATs decreases thermotolerance

Cultivar-specific gene regulation is critical to delineate regulatory networks involved in tolerance mechanisms under varied environmental conditions (Sun et al., 2010; Koeslin-Findeklee 2014; Boccacci et al., 2017; Balyan et al., 2017). This study identifies 21 antagonistcally expressing genes upon HS in the two contrasting varieties, the differential expression of 10 of which was also corroborated by qRT-PCR (Figure 2D). Surprisingly, the direct role of these genes in hyperthermal stress tolerance is either totally lacking or very limited. Therefore, to establish the regulatory influence of these genes in governing thermotolerance, we performed *in-planta* functional analysis using overexpression and virus induced gene silencing (VIGS) in tomato. To short-list few candidate genes for this analyses, we put another filter by assessing the transcript abundance of the above 10 genes under HS in another pair of contrasting cultivars namely, Pusa Sadabahar from the tolerant clade and Pusa Ruby from the sensitive clade (Supplementary Figure S11 and Figure 1). We selected two genes (Solyc09g014280 and Solyc07g056570) that are up-regulated in CLN as well as Pusa Sadabahar while are down-regulated in both sensitive cultivars (‘tolerant-up genes’) as well as one gene (Solyc03g020030) which is highly down-regulated in tolerant CLN but up-regulated in both sensitive cultivars (‘sensitive-up gene’).

*Solyc09g014280* gene codes for an *HXXXD-type acyl-transferase* and belongs to the acylsugar acyltransferases (ASATs) class ofthe large and diverse BAHD-family of acyltransferases (Moghe et al., 2017). Acylsugars are a group of small specialized metabolites having diverse structures that are restricted to plants in the Solanaceae family that act as natural chemicals against pests like whiteflies and spider mites (Alba et al., 2009; Liedl et al., 1995). Acylsugars are produced in the tip cell of the long glandular secreting trichomes using ASAT enzymes that catalyse sequential addition of specific acyl chains to the sucrose molecule using acyl CoA donors (Schilmiller et al., 2008; Maldonado et al., 2006). *Solyc09g014280 and Solyc05g052670* (another tolerant-up gene but not being functionally characterised, Supplementary Figure S11) are reported to be enriched in the glandular trichomes in tomato (Moghe et al., 2017). While *Solyc09g014280* is induced upon potato cyst nematode infection in tomato roots (Swiecicka et al., 2017), *Solyc05g052670* is up-regulated in tomato in response to tomato yellow leaf curl virus infection (Chen et al., 2013). Here we report yet unexplored function of an ASAT member in tomato HS response. Plants exhibiting successful knock-down of *Solyc09g014280* transcripts (Supplementary Figure S12) showed severe drooping of leaves upon recovery after being exposed to HS in comparison to vector-control plants (Figure 4A). Heat stress results in a higher than optimal concentration of ROS that not only affect plant’s ability to photosynthesize but also causes cell death. These parameters are routinely adopted to assess thermotolerance of plants. The suppression of ASAT gene shows enhanced hydrogen peroxide (one of several reactive oxygen species) levels as well as cell death under HS as judged by DAB staining and Trypan blue staining, respectively (Figure 4B). Gas exchange parameters measured by Li-Cor 6400 portable photosynthesis measuring system at the end of the heat treatment of *ASAT* silenced gene showed significant drop in net photosynthesis rate and water use efficiency (WUEi) as well as enhanced stomatal conductance and transpiration rate (Figure 4C-F) highlighting the role of ASAT as a positive regulator of thermotolerance. This conclusion was further strengthened by assessing seedling performance under HS when *ASAT* gene was over-expressed. The seedling survival increased by nearly 17% in comparison to vector control (Figure 4G) and the reduction in hypocotyl length under HS was also around 8% in overexpression plants in comparison to 18% in vector control plants (Figure 4H). Moreover, the relative abundance of HS marker genes (HSFA2a/A7a/A3a, sHSP24.5 CI and HSP90) (Figure 4I) was reduced several folds in *ASAT* silenced plants reiterating the fact that ASAT has a key role in providing thermotolerance. Further, to understand how the transcriptional regulation of *ASAT* results in antagonistic expression between the contrasting cultivars, we sequenced 1.5kb promoter from both the cultivars and checked promoter ASAT:GUS reporter expression by assaying CLN promoter: GUS-reporter in CA4 background and vice-versa. We find that there is one substitution of ‘T’ to ‘A’ in CA4 promoter, which disrupts the binding sites of two TFs namely, NFY-A/B/C and AP2 having *cis*-binding sites i.e. (ACAAT) and(AA<u>T</u>CAA) respectively, in CA4. If this mutated regulatory site abrogates TF binding and in turn *ASAT* expression in CA4 then the expression of CA4-ASAT promoter:GUS in CLN background is expected to reduce, indeed this was the case (Figure 4J). However, since the expression of CLN-ASAT promoter:GUS is up-regulated in CLN background (Figure 4J), but not in CA4 background, it appears that in addition to the promoter sequence context, cultivar-differential TF pool between CLN and CA4 also regulate the expression of this gene. *ASAT* promoter has binding sites for more than 15 different TFs (Supplementary Figure S13). Many of these TFs showed highly differential expression between the two cultivars as well as cultivar-biased expression (Supplementary Table S6 and Figure 3D). Our data provides evidence of a novel role to ASAT as a positive regulator of tomato thermotolerance. The exact mode of action of this enzyme under HS needs further attention.

**Figure 4.**
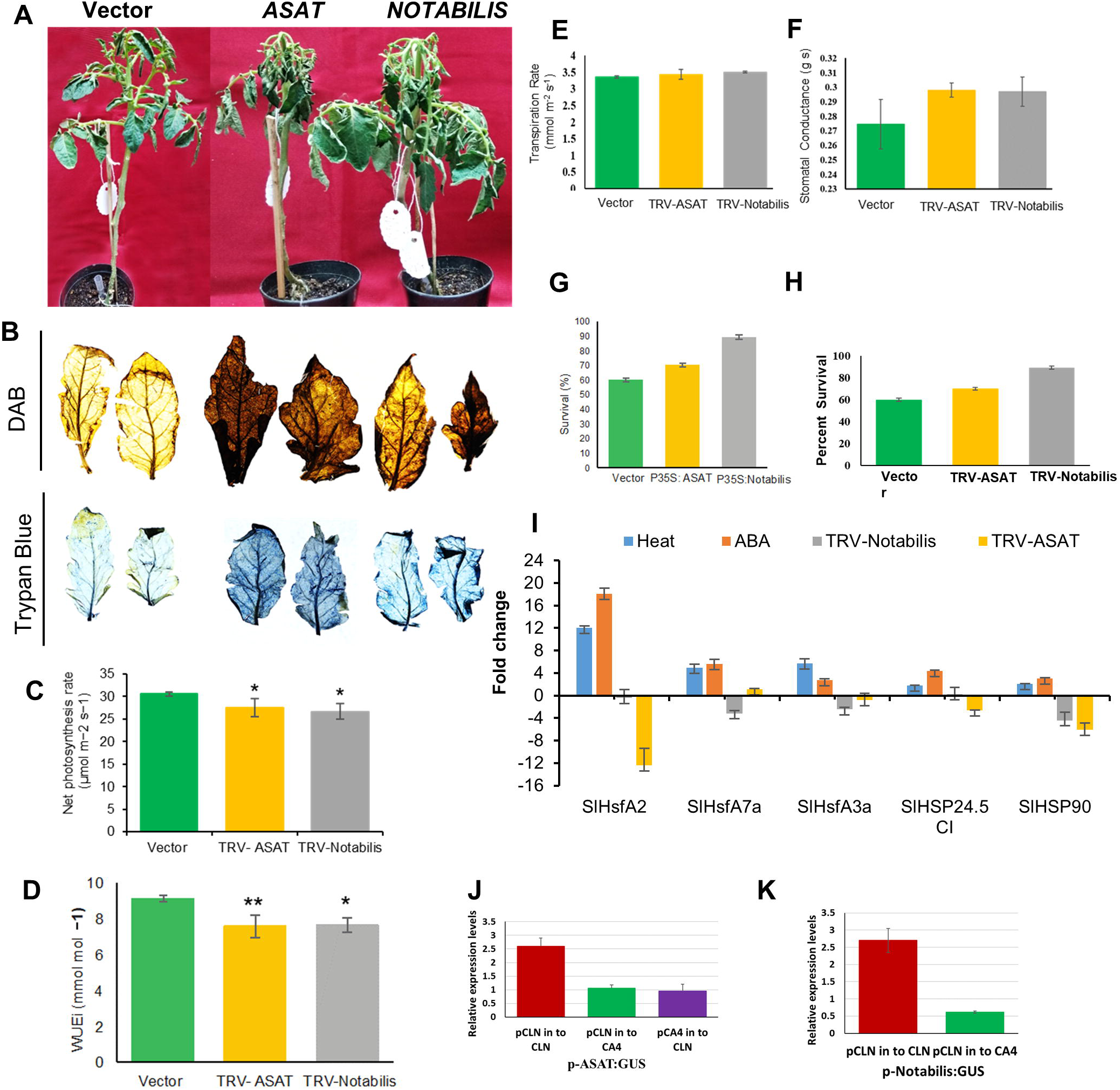
Functional validation of the roles of *ASAT* and *Notabilis* genes in response to heat stress. **(A)** Phenotypes of HS treated empty vector (TRV), TRV-ASAT and TRV-Notabilis silenced plants. 15-days-old CLN plants were agro-infiltrated with empty vector (TRV), TRV-ASAT and TRV-Notabilis VIGS constructs. Plants were assayed for heat stress tolerance after 3 weeks of infiltration. **(B)** DAB and trypan blue staining of leaves of HS treated empty vector (TRV), TRV-ASAT and TRV-Notabilis silenced plants. **(C-F)** Estimation of net photosynthesis rate (µmol m^−2^s^−1^), water use efficiency (mmol mol^−1^), stomatal conductance (mol m^−2^s^−1^), and transpiration rate (mmol m^−2^s^−1^) in empty vector (TRV), TRV-ASAT and *TRV-Notabilis* silenced plants following heat stress. **(G-H)** Estimation of survival (percent) and % reduction of hypocotyl length of seedlings under control conditions and after 5 days of heat stress in empty vector (TRV), *TRV-ASA T* and *TRV-Notabilis* overexpressing seedlings. Data are means of four biological sets of 70 seedling each. **(I)** Expression profiles of HSR genes in heat, ABA and VIGS silenced *ASAT* and *Notabilis* plants by qRT-PCR. **(J)** GUS:reporter assays of CLN and CA4 ASAT promoters in CLN and CA4 background. **(K)** GUS:reporter assays of CLN:Notabilis promoter in CLN and CA4 background. Error bars denote standard error. *P<0.05, **P<0.01 and ***P<0.001

### Silencing Notabilis confers thermosensitivity in tomato

We then functionally validated the other tolerant-up gene, *Solyc07g056570* or *Notabilis* (9*-cis-epoxycarotenoid dioxygenase 1*). Notabilis regulates the rate limiting step in the ABA biosynthesis pathway, and has important role in water stress tolerance as evidenced by several reports (Burbidge et al., 1999; Qin and Zeevaart, 1999; Luchi et al., 2001; Wan and Li, 2006; He et al., 2018). In Arabidopsis, ABA deficient and signalling mutants are involved in acquired thermotolerance acquisition (Larkindale et al., 2004). In lettuce, its homolog (*NCED4*) is the causal gene in the Htg6.1 QTL, associated with thermo-inhibition of seed germination (Huo et al., 2013). Here we investigate the role of Notabilis in tomato thermotolerance. When *Notabilis* knocked-down plants (Supplementary Figure S12) were exposed to HS and assessed after recovery, they exhibited decreased thermotolerance as evidenced by the morphological (severe wilting of leaves) phenotype (Figure 4A). This is in accordance with the phenotype observed in the tomato null mutant of *notabilis* that is deficient in ABA synthesis (Burbidge et al., 1999; Thompson et al., 2000, 2004). It is known that high ABA levels cause ROS outburst. However, the leaves of *Notabilis* silenced plants show slightly increased ROS levels and HS-induced cell death in comparison to vector-control plants (Figure 4B). This apparent anomaly may be attributed to a balance between low ROS levels due to reduced ABA levels and other pathways contributing to ROS production in response to HS. The gas exchange parameters analysis further supported the important role of Notabilis in thermotolerance. There was significant reduction in the photosynthetic rate and water use efficiency along with a significant rise in stomatal conductance and enhanced transpiration rate in silenced plants in contrast to vector control plants (Figure 4C-F). Further proof for *Notabilis* as agene imparting thermotolerance was provided from tomato plants over-expressing *Notabilis* as judged by better seedling survival percentage and hypocotyl length in comparison to vector control (Figure 4G-H). We then measured the relative abundance of HS marker genes in the silenced *Notabilis* plants; the level of all the genes was reduced several folds upon HS in comparison to control (Figure 4I). Moreover, when we checked the expression of these HSR genes in response to ABA, they were all up-regulated suggesting *Notabilis* as an upstream regulator to these HS marker genes(Figure 4I). Of these HSFs, HSFA2a and HSFA7a were down-regulated the most and appeared as the critical candidates in the Notabilis-ABA pathway mediated thermotolerance. HSFA2a is well known heat stress regulator (Scharf et al., 2012) we therefore,chose to assess HSFA7a forthermotolerance. Indeed, over-expression of HSFA7a made tomato seedlings more HS tolerant as measured by seedling survival, hypocotyl length and seedling weight (Supplementary Figure S14). Further, we asked whether the opposite expression of *Notabilis* in CLN-CA4 cultivars is because of variation in promoter sequences and/or an inherent difference in TF pool of both the cultivars. Sequence information of promoters for both the varieties highlighted no SNPs, thus a regulatory influence of the *cis*-sites in the contrasting expression was ruled out. We then checked the expression of GUS gene hooked to *Notabilis* promoter in CLN and CA4 background. Figure 4K highlights that GUS expression is up-regulated only in CLN background, pointing to a probable cultivar-specific TF-mediated regulation. In literature, *Notabilis* is shown to be transcriptionally regulated by SlNAP2 (Solyc04g005610) (Ma et al., 2018) a NAC TF. Indeed, this TF is up-regulated in CLN (~2-folds) but not in CA4 in response to HS (Supplementary Table S3).

### Silencing PI-II gene enhances tomato thermotolerance

Further, another gene but a ‘sensitive-up gene’ which is *Solyc03g020030* was functionally characterised. This belongs to the proteinase inhibitor-II (PI-II) serine-PI family having dimeric domains in the protein, inhibiting trypsin and chymotrypsin (Tamhane et al., 2012). The PI-IIs are well documented in plant defense against insects, bacteria and fungi (Rehman et al., 2017). Few reports in abiotic stress tolerance like drought, salinity, osmotic variations and pH are also documented (Huang et al., 2007; Shan et al., 2008; Srinivasan et al., 2009). Only a Kunitz PI (another class of serine-PI family) has been shown to be induced in response to heat stress (Annamalai and Yanagihara 1999) however, information regarding PI-II-class role in HS is completely lacking. Solyc03g020030 is expressed at high levels in CLN under control condition but decreasesup to 3-folds under HS while significant increase was noticed in CA4 under HS. We find, there is enrichment of GO term of ‘serine-type endopeptidase inhibitor activity’ in CA4 specific up-regulation (Figure 2C). Silencing of *PI-II* was confirmed by qRT-PCR (Supplementary Figure S12) and the leaves of knock-down plants were upright, enduring HS better than those of vector-control plants which showed drooping and burnt leaves (Figure 5A). Moreover, there was appreciable difference phenotypically when VIGS was repeated in sensitive background (Pusa Ruby) (Figure5B). The CLN *PI-II* knockdown plants exhibited significant enhancement in WUEi and net photosynthetic rates as compared to TRV-control plants under HS (Figure 5C). In addition, the transpiration rate and the stomatal conductance were lower in TRV-PI-II plants under HS (Figure 5C-D). The *PI-II* overexpression in CLN decreased the tolerance of this robust cultivar’s seedlings which exhibited much lower hypocotyl length and survival under HS as compared to vector control seedlings. In contrast VIGS based silencing of of *PI-II* in pusa ruby exhibit higher survival rate (Figure 5E-G). Further, the relative abundance of different HS marker genes in the silenced *PI-II* plants enhanced upon HS in comparison to control confirming its role as a negative regulator of HS-tolerance (Figure 5H-I). In tomato, *PI-II* is transcriptionally regulated by SlbZIP1 TF (Solyc01g079480.3) (Zhu et al., 2018). When we checked our NGS data for a possible cultivar-specific differential response of *PI-II* by SlbZIP1in response to HS, the TF was not significantly differentially regulated in CLN (1.85-folds change) and CA4 (1.25-folds change) (Supplementary Table S3). This suggested that HS-mediated regulation of *PI-II* is governed by other TFs or has epigenetic regulation. The down-regulation of *PI-II* by VIGS established its role in retrograde signalling in the regulation of HS. Knocking-down this gene by CRISPR/RNAi technology could be highly promising for engineering for HS-thermotolerance.

**Figure 5.**
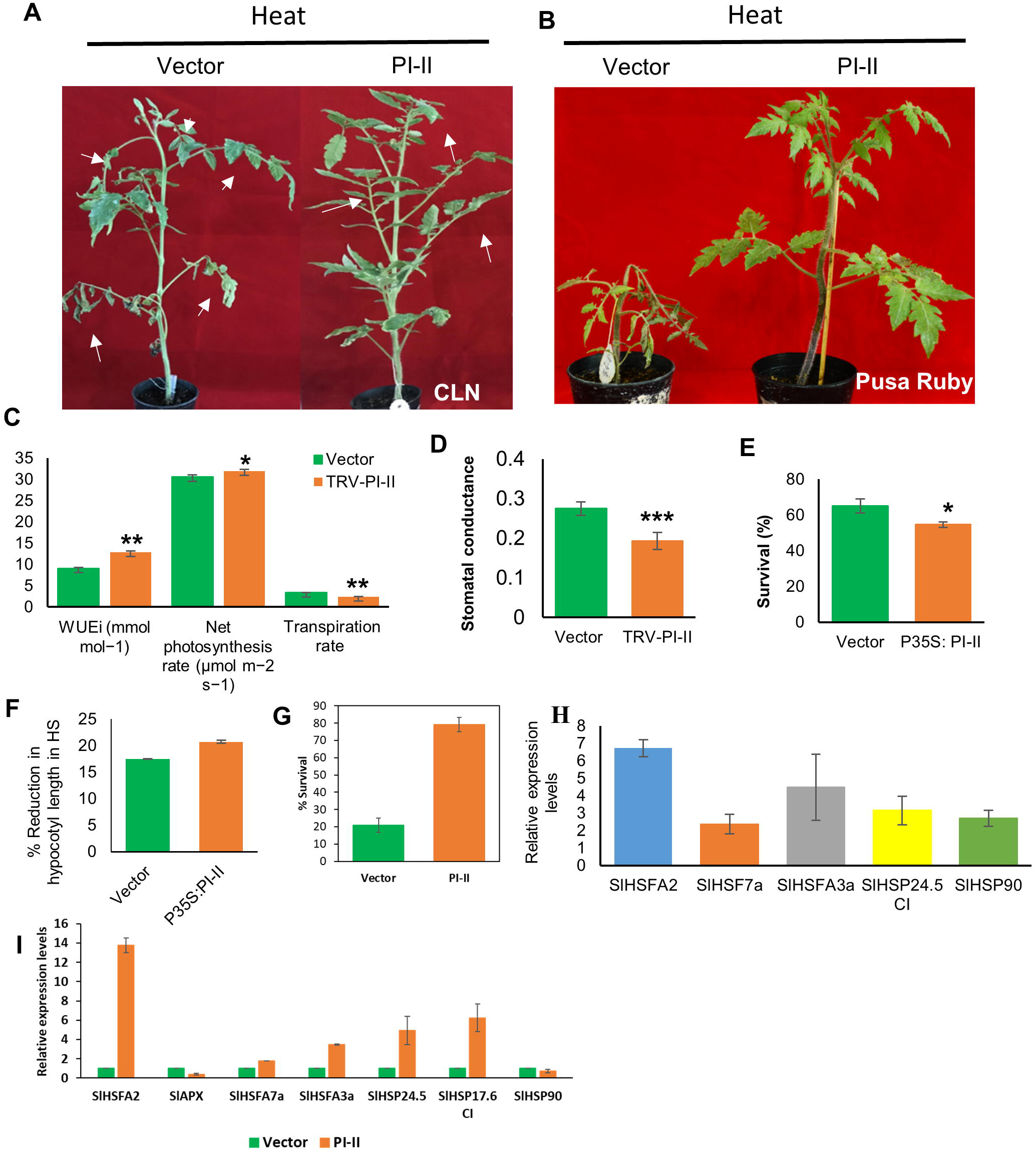
Functional validation of Pin-II type proteinase inhibitor in response to HS. **(A)** 15 days old CLN plants were agro-infiltrated with empty vector (TRV) and TRV- pin-II type proteinase inhibitor VIGS construct. Plants were assayed for HS tolerance after 3 weeks of infiltration. Phenotypes of HS treated TRV and TRV-pin-II type proteinase inhibitor plants was assessed in tolerant cultivar CLN (A) and in sensitive cultivar Pusa Ruby**(C)** Estimation of water use efficiency (mmol mol^−1^), net photosynthesis rate (µmol m^−2^s^−1^), transpiration rate (mmol m^−2^s^−1^) and **(D)** stomatal conductance (mol m^−2^s^−1^) in TRV and TRV-pin-II type proteinase inhibitor plants of CLN following heat stress. **(E)** Estimation of seedling survival (percent) and **(F)** % reduction in hypocotyl length during HS in vector control and Pin-II type proteinase inhibitor overexpression seedlings. **(G)** Estimation of survival rate in empty vector (TRV) and TRV-pin-II type proteinase inhibitor VIGS silenced plants in Pusa ruby after heat stress. **(H-I)** Expression profiles of HSR genes in VIGS silenced Pl-II plants in tolerant cultivar CLN (H) and in sensitive cultivar Pusa Ruby (I). Data are means and SE of four biological sets of 70 seedling each. *P<0.05, **P<0.01 and ***P<0.001

## Conclusions

This study has identified a contrasting pair of tomato cultivar showing heat stress tolerance and sensitivity from varieties growing in the sub-tropical Indian climatic zones. Further, we have utilized their comparative transcriptome to identify many novel heat stress regulators which have not been examined previously in thermotolerance. The efficacy of their significance in enhancing thermotolerance has been demonstrated for three transcription factors, two genes reported so far only in plant defense and a key enzyme in ABA hormone biosynthesis. In addition, many promising genes have been identified which can serve as excellent candidates for enhancing thermotolerance in tomato and be assessed and used in other crop plants. Their utility in providing robustness to plants in other abiotic stresses needs to be evaluated.

## Large datasets

## Acknowledgments

This work is supported by grants from NIPGR and SERB (EMR/2016/006229). The authors acknowledge NIPGR phytotron facility, CIF and field area. CLN1621L and CA4 seeds were provided by AVRDC, Taiwan. SR acknowledges Department of Biotechnology (DBT) Govt. of India, SJ and CB acknowledge University Grants Commission (UGC) and JRD acknowledges Council of Scientific and Industrial Research (CSIR) for the award of research fellowships.

## Supplementary Figure legends

**Supplementary Figure S1**. Average day/night temperature (°**C**) in field for tomato cultivar assessment for thermotolerance. Data recorded in year 2014-2016 during weeks of tomato cultivars plantation in experimental fields of National Institute of Plant Genome Research (NIPGR), New Delhi, India. (A) Normal Season average temperature (°**C**) (Week-1; 4th week of October 2014 and 2015). (B) Warm season average temperature (°**C**) (Week-1; 4th week of February 2015 and 2016). (C) The average day/night temperature (°**C**) during the flowering in normal season and warm season. Black Arrow in ‘A’ and ‘B’; start time duration of flowering in tomato cultivars in both seasons.

**Supplementary Figure S2.** Comparative analysis of various tomato cultivars in response to HS. (A-B) The survival assays performed at 5-days-old seedlings (A) and 1-month-old plants (B) in response to heat stress. (C-E) The measurement of proline content, relative water content and electrolytic leakage in leaves of 1-month-old plants of different cultivars under heat stress. The heat regime for A was basal heat stress at 45°C for 4.5 h and for B–E was gradual increase in temperature from 26 °C to 45 °C for 4h and then plants were exposed for 4.5 h at 45°C. The effect of heat stress on percent fruit set and tomato yield of different cultivars (F-G). The assessment was done over 2 years with data collected in normal growing season (November to February; 2014-2015 and 2015-2016) and warm season (March to June; 2015 and 2016). The error bars depict standard error. *p<0.05, **p<0.01 and ***p<0.001.

**Supplementary Figure S3. Transcriptome analysis of CLN and CA4 leaf under control and heat stress. (A)** PCA plots of all RNA-seq libraries of CLN and CA4 under control and heat stress regimens. **(B and C)** Hierarchical clustering of different samples **(B)** and genes **(c)** in response to heat stress in CLN and CA4. The heat maps were plotted using the log_2_ transformed RPKM values following the average clustering algorithm using CLC Genomics Workbench software. **(D)** Volcano plots of all the expressed genes in CLN-C vs. CLN-H and CA4-C vs. CA4-H wherein the X and Y axis represents the fold change and –log_10_ (FDR p-value), respectively.

**Supplementary Figure S4. The GO term enrichment analysis of gene set showing conserved up-regulation in response to heat stress.** The table showing enrichment analysis for biological process (A), molecular function (B) and protein class (C) (fold enrichment ≥2 and p-value ≤0.05) in the genes showing up-regulation in both (CLN and CA4) in response to heat stress.

**Supplementary Figure S5. The GO term enrichment analysis of gene set showing conserved down-regulation in response to heat stress.** The table showing enrichment analysis for biological process (fold enrichment ≥2 and p-value ≤0.05) in the genes showing down-regulation in both (CLN and CA4) in response to heat stress

**Supplementary Figure S6. The GO term enrichment analysis of gene set showing conserved down-regulation in response to heat stress.** The table showing enrichment analysis for molecular function (fold enrichment ≥2 and p-value ≤0.05) in the genes showing down-regulation in both (CLN and CA4) in response to heat stress.

**Supplementary Figure S7. The GO term enrichment analysis of gene set showing conserved down-regulation in response to heat stress.** The tables showing enrichment analysis for cellular component (A) and protein class (B) (fold enrichment ≥2 and p-value ≤0.05) in the genes showing down-regulation in both (CLN and CA4) in response to heat stress.

**Supplementary Figure S8. The GO term enrichment analysis of gene set showing tolerant cultivar specific HS response.** The tables showing enrichment analysis for biological process (A), molecular component (B) and cellular component (C) for genes exhibiting downregulation in response to HS only in CLN and not in CA4. The genes with significant down (≥2 FC and p-value ≤0.05 in CLN; ≤2 FC in CA4) were considered.

**Supplementary Figure S9. The GO term enrichment analysis of gene set showing sensitive cultivar specific upregulation.** The tables showing enrichment analysis for biological process (A), molecular component (B) and protein class (C) for genes exhibiting upregulation in response to HS only in CA4 and not in CLN.

**Supplementary Figure S10. The GO term enrichment analysis of gene set showing sensitive cultivar specific downregulation.** The tables showing enrichment analysis for biological process (A), molecular component (B) and protein class (C) for genes exhibiting downregulation in response to HS only in CA4 and not in CLN.

**Supplementary Figure S11. Expression analysis of antagonistically selected genes in different tolerant and sensitive cultivars.** The qRT-PCR expression analysis of genes with inverse HS-response in contrasting cultivars in leaf of heat tolerant (CLN and PusaSadabahar) and sensitive (CA4 and Pusa Ruby) under heat stress conditions. For each condition two biological and three technical repeats were used. The expression levels of genes were calculated using the 2−ΔΔCt method and presented using fold change values transformed to log_2_ relative to respective Actin gene expression. The error bars represent standard error.

**Supplementary Figure S12. Virus Induced Gene Silencing of *Acylsugaracyltransferase, Notabilis* and *Pin-II type proteinase inhibitor 1 (PI-II).*** Quantitative RT-PCR analysis of *ASAT, Notabilis*and PI-II genes in TRV and TRV-ASAT, TRV-*Notabilis* and TRV-*PI-II* plants confirming the silencing of their respective genes. The expression levels of genes were calculated using the 2−ΔΔCt method and presented using fold-change values transformed to log 2 format compared with control.

**Supplementary Figure S13. TF families and cis-elements associated with *Acylsugaracyltransferase (ASAT)* promoter. (A)** The number of various binding sites (cis-elements) of different transcription factors in the promoter of ASAT as predicted by PlantPAN database

**Supplementary Figure S14. Functional validation of the role of HSFA7a in heat stress response. (A)** Five-days-old CLN seedlings were agro-infiltrated with overexpression construct of HSFA7a, plants were assayed for HS tolerance after 2 days of infiltration. Phenotypes of HS treated plants were analysed for survival and hypocotyl length after 6 days of recovery. **(B)** Phenotype of HSFA7a overexpressing and vector control seedlings after HS **(C)** Estimation of survival rate of HSFA7a overexpressing and vector control seedlings following heat stress, hypocotyl length and 30 seedlings dry weight under HS after 6 days of recovery in vector control and HSFA7a overexpression seedlings. Data are means of four biological sets of 70 seedlings each. The error bars depict standard error. *p<0.05, **p<0.01 and ***p<0.001.

## Supplemenatry Table legends

**Supplementary Table S1**List of primers used in the study.

**Supplementary Table S2: Summary of paired-end dataset analysis**. The ITAG version 3.2 gene model at SGN (Sol Genomics Networks) was used as the reference for mapping.

**Supplementary Table S3:** The list and expression level (RPKM and fold change) of all the genes in the study. The expression level (RPKM) and fold change of all the genes (35,768) annotated in ITAG version 3.2 in CLN and CA4 under control and heat stress conditions.

**Supplemental table S4.** KEGG pathway enrichment analysis of HS responsive genes in leaf of CLN and CA4.

**Supplementary Table S5:** The list of genes with cultivar specific expression under control and HS condition.

**Supplementary Table S6:** The expression level of transcription factors in leaves of CLN and CA4 under control and HS conditions.

**Supplementary Table S7**: HS regulation of heat shock proteins in CLN and CA4

**Supplemenatry methods S1**

